# OPTN limits herpes stromal keratitis severity and demyelination through negative regulation of IL-17 and hyperinflammatory T-cell response

**DOI:** 10.1101/2021.11.29.470418

**Authors:** Joshua Ames, Tejabjiram Yadavalli, Chandrashekhar Patil, James Hopkins, Ilina Bhattacharya, Deepak Shukla

## Abstract

Herpes stromal keratitis (HSK) is a result of the inflammatory sequelae following primary and recurrent Herpes simplex virus type-1 (HSV-1) infections. This pathology is known to be mediated by immunopathogenic T cell responses against viral antigens, however most individuals infected with HSV-1 never exhibit signs of this immunopathology. Recent studies have identified the host restriction factor, optineurin (OPTN), as an inhibitor of viral spread in the central nervous system, protecting hosts from viral encephalopathy. In an HSV-1 corneal infection mouse model on OPTN knockout mice, we assess the contribution of OPTN to ameliorating the clinical manifestations of HSK. We identify that OPTN protects the host from loss of ocular and whisker sensitivity and opacification of the cornea. scRNA-seq of the trigeminal ganglion (TG) reveals that transcription changes to the peripheral neurons and immune cell populations drive the expression of Il-17A in CD4 and CD8 T cells, as well as increased infiltration of T cells into the TG. This leads to demyelination and the observed HSK pathology.

## Introduction

Upon HSV-1 infection, the virus travels to the peripheral neurons of the trigeminal ganglion (TG) where it establishes lifelong latency ^1^. In most individuals the virus and associated inflammation is suppressed indefinitely, but in some cases the complications are caused by recurrent reactivation events ^2^. Herpes stromal keratitis (HSK) is a degenerative condition caused by immune responses to recurrent herpes virus infections of the cornea and it presents clinically as inflammation and scarring of the cornea often with the presence of dendritic lesions and diminished sensitivity of the cornea ^3^. This is a result of reactivation of latent herpes infection, and only long-term treatment with antiviral and steroids can halt progression to corneal blindness ^4^. Herpes simplex virus type-1 (HSV-1) is the most common cause of HSK; however it is poorly understood why most individuals can control the infection and why there is often a disconnect between viral shedding and clinical presentation.

We recently identified OPTN as a host intrinsic restriction factor that protects against neuroinvasive herpes infections of the central nervous system. Through selective autophagy of HSV-1 proteins and suppression of RIPK1 dependent cell death, OPTN acts as an intrinsic immune barrier in neurons of the central nervous system (CNS) ^5^. While this explains the antiviral role of OPTN in the CNS, we are interested in the role of OPTN in the more common orofacial and ocular presentations of herpes infection. Herein we show that OPTN suppresses hyperinflammatory responses independently of the level of infection, and this results in severe herpes keratitis and damage to TG.

## OPTN restricts severity of Herpes Stromal Keratitis

To study the role of OPTN in herpes keratitis, we use an *Optn*^−/−^ mouse model where animals were unilaterally infected by application of HSV-1 to the cornea following scarification. *Optn*^−/−^ animals showed a rapid and permanent loss of corneal sensitivity by 4 dpi that was still present 26 dpi while the *Optn*^+/+^ did not lose much, if any, sensitivity (figure 1A). Unexpectedly, we found that the *Optn*^−/−^ animals had also lost bilateral whisker sensitivity by 26 dpi (figure 1B). Scarring due to HSK can lead to opacification of the cornea, and there was significantly more severe corneal opacification in the *Optn*^−/−^ animals which resulted in a nearly completely opaque cornea by 8 dpi (figure 1C-D).

**Figure 1:**
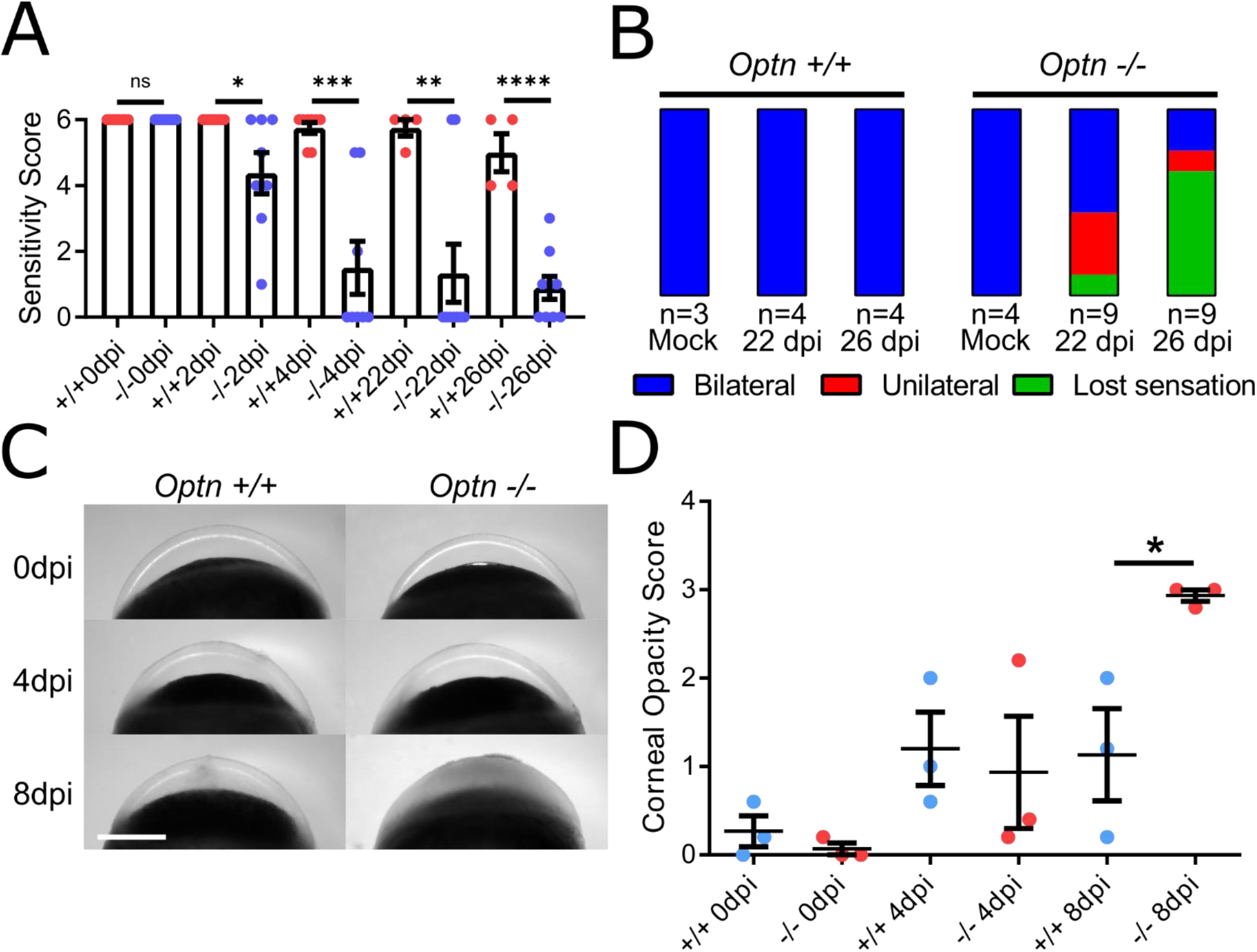
*Optn* knockout increases Herpes stromal keratitis severity and leads to lost corneal and facial sensation. A) Wildtype or *Optn* knockout mice were infected with 1×10^5^ PFU HSV-1 then A) corneal sensitivity was measure by blink response at 0, 2, 4, 22, and 26 dpi and B) Whisker sensitivity is reported as bilateral (full sensitivity), unilateral (sensitivity lost on one side of muzzle), or lost sensation. n >/= 3 mice per group. n >/= 3 mice per group. C) Shown are representative photographs of mouse corneas, highlight the opacification with infection. Scale bar is 1 mm. D) Quantification of mouse corneal opacity scores following infection. n = 3 mice per group. Student’s t-test was performed for statistical analysis (α= 0.05). *p < 0.05; **p < 0.01; ***p < 0.001, ****p < 0.0001, ns, not significant.

Despite these differences in herpes keratitis, tissue titers at 4 dpi or 8 dpi and ocular wash titer at 2 dpi and 4 dpi revealed no significant difference in the levels of HSV-1 replication between the two mouse genotypes (figure 2A-B). Histology shows that at 4 dpi and 8 dpi both the *Optn*^+/+^ and *Optn*^−/−^ mice show similar levels of inflammation and corneal thickening (figure 2C). These results suggest that OPTN protects primarily against lost of sensation of the eyes and whiskers and may be protecting the trigeminal ganglion and supporting neurons.

**Figure 2:**
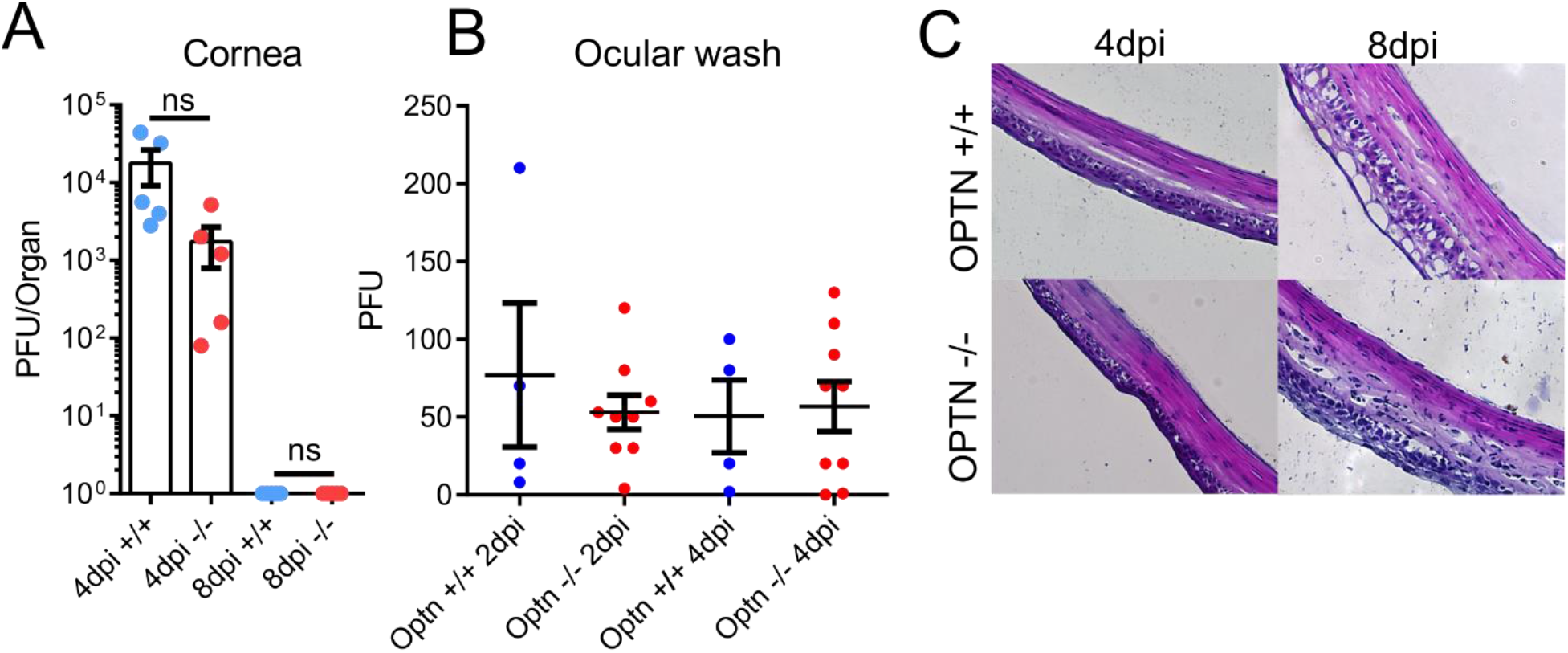
*Optn* knockout does not alter HSV-1 replication in the mouse eye. A) Wildtype or *Optn* knockout mice were infected with 1×10^5^ PFU HSV-1 and corneal tissue HSV-1 titers were measure at 4 and 8 dpi. B) Ocular wash HSV-1 titers from 2 and 4 dpi were measured. C) Representative H&E staining of HSV-1 infected corneas reveal similar levels of inflammation. Student’s t-test was performed for statistical analysis (α= 0.05). ns, not significant.

## scRNA-seq of 30 dpi trigeminal ganglion reveal differences between *Optn*^+/+^ and *Optn*^−/−^ mice

After observing the increased ocular pathology in *Optn^−/−^* mice without a commensurate increase in HSV-1 replication in the cornea and detecting the permanent loss of corneal and whisker sensitivity in the *Optn*^−/−^ mice, we hypothesized that there must be permanent changes in the TG of the *Optn*^−/−^ mice that lead to peripheral neurodegeneration. We performed DROPseq to generate single-cell cDNA libraries of cells prepared from 30 dpi *Optn*^+/+^ or *Optn*^−/−^ mouse TG where three TG were pooled per treatment. We chose a reduced model to highlight difference only during the chronically diseased or recovered state. These were then sequenced, annotated, and analyzed for differential gene expression (DGE) and pathway enrichment.

We sequenced a total of 2746 cells. Broad cell type identities were assigned using literature-based annotation. We were able to identify the major cell types of the TG, including peripheral neurons (pNeurons), myelinating Schwann cells (mSchwann), non-myelinating Schwann cells (nmSchwann), macrophages, endothelial cells, and epithelial cells (figure 3A-C). It was immediately apparent in a UMAP dimensionality reduction analysis that the *Optn*^−/−^ have much lower proportion of peripheral neurons sequenced compared to the *Optn*^+/+^ sample. Additionally, Pathway analysis of differentially expressed gene between *Optn*^+/+^ and *Optn*^−/−^ within a cell type revealed multiple differentially expressed pathways across cell types with the most enriched pathways represented in the pNeuron, Macrophage, Endothelial, and Lymphoid cell types (figure 3D). This loss of neuronal activity and differential expression may underlie the *in vivo* loss of sensation observed.

**Figure 3:**
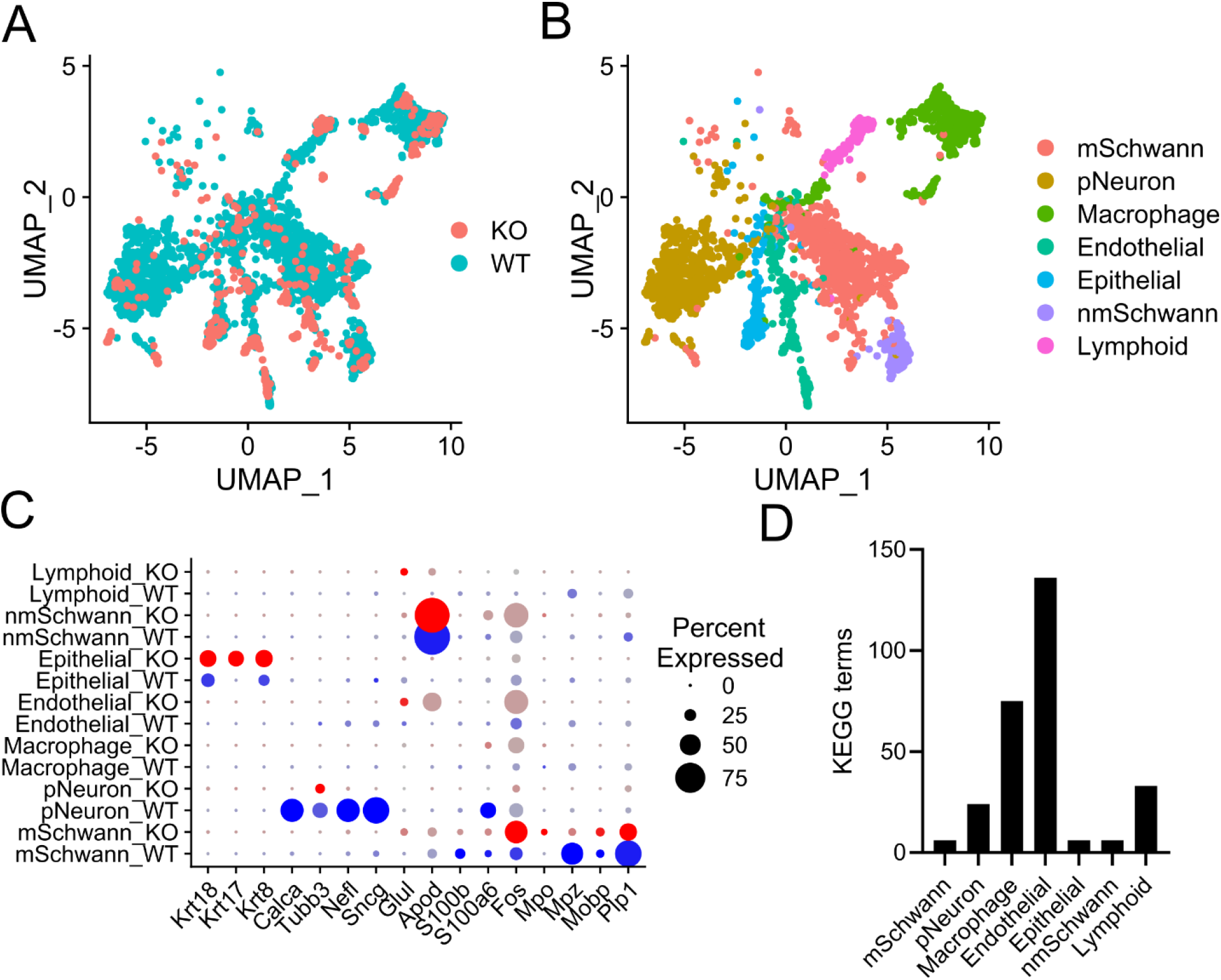
scRNAseq of 30 dpi trigeminal ganglion reveal differences between WT and KO mice. scRNA-seq was performed on 3 pooled mouse trigeminal ganglion at 30 dpi. A) UMAP analysis reveals the distribution of cells from *Optn^+/+^* (WT) and *Optn*^−/−^ (KO) across clusters in the dataset. B) UMAP analyses annotated with putative cell types based on highly represented cluster biomarkers. C) Dot plot highlighting the expression of several cell type markers by cell type and genotype. D) Graph of the number of KEGG pathways detected in an enrichment analysis using the differentially expressed genes between KO and WT cells within each cluster.

## Single cell analysis reveals differences in immune expression signatures and differentially regulated pathways

Immune markers that there highly represented in *Optn*^−/−^ cell types are consistent with observation of chronic inflammatory symptoms including keratitis and loss of sensation (figure 4A). Both *Optn*^+/+^ and *Optn*^−/−^ macrophage populations expressed immune effector genes including Lyz1/2, C1qb. C1qc, C3, Ly86, and MHCII genes, H2-Aa and H2-Ab1. Only *Optn*^−/−^ macrophages show high expression of Ccr2, Ccl4, Ly6a2 and Ly6c2. Endothelial cells show increases in Ifit1, Ifit2, Ifit3, Cxcl9, Cxcl1, Icam1, and Vcam1, which is consistent with recruitment of circulating lymphocytes to the trigeminal ganglion. *Optn*^+/+^ Lymphoid cells show increased expression of several markers compared to *Optn*^+/+^ Lymphoid cells including Cd69, Cd28, Cd8b1, and Cd8a, consistent with Cd8 T cell recruitment. Furthermore, *Optn*^−/−^ lymphoid cells have higher levels of Il17ra, Il7r, Il2rg, Cxcr6, and Cxcr3 which are markers for infiltrating inflammatory Cd8 T cells.

**Figure 4:**
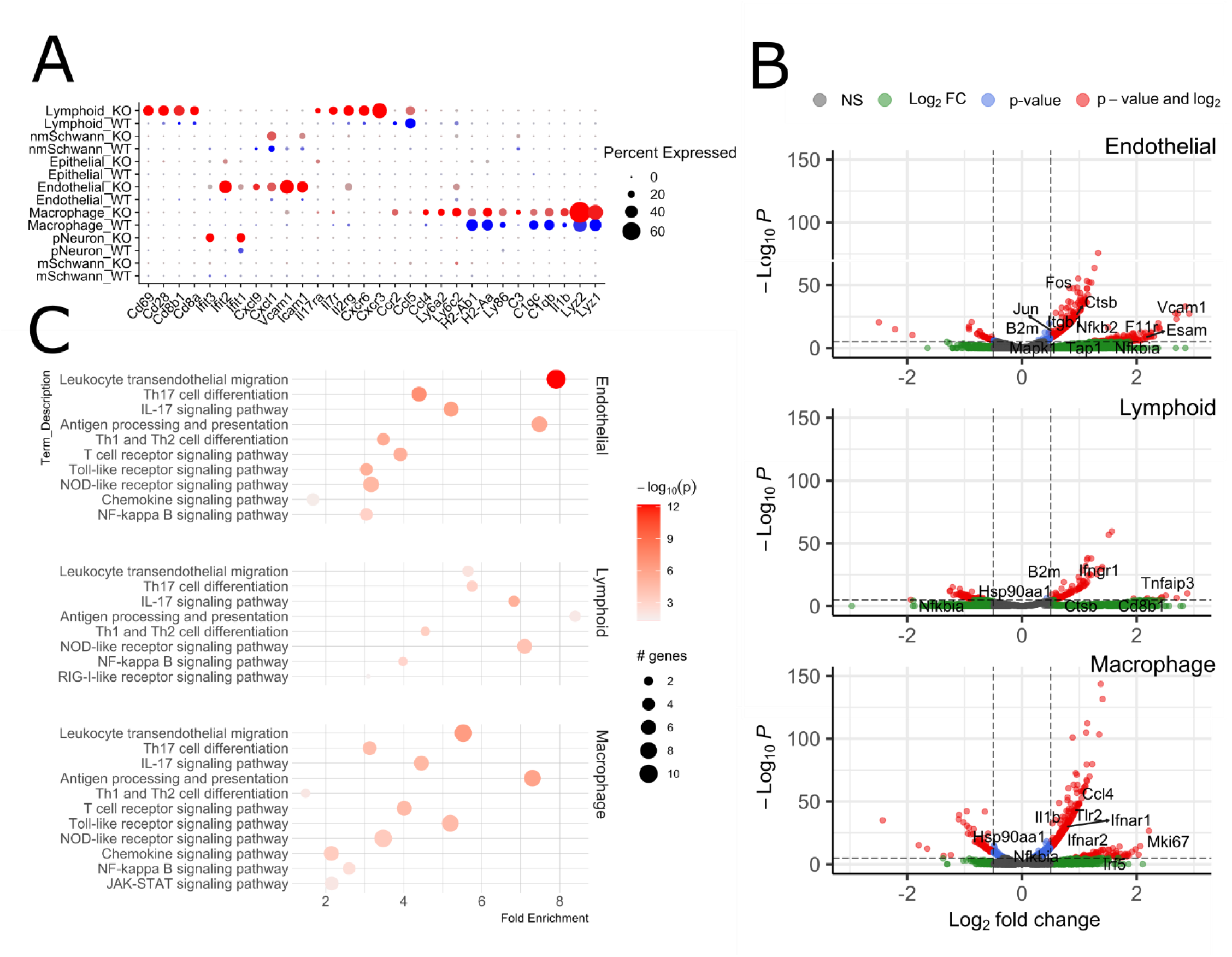
single cell analysis reveals differences in immune expression signatures and differentially regulated pathways. A) Dot plot highlighting the expression of several immune system related markers by cell type and genotype. B) Volcano plots of differentially expressed genes in Lymphoid, Macrophage, or Endothelial cell types. C) Dot plot of significantly enriched immune related pathways detected in Lymphoid, Macrophage, or Endothelial cell types.

The observed increase in markers across cell types that would support increased inflammation in the *Optn*^−/−^ TG was supported by differential gene expression and KEGG pathway enrichment analysis (figure 4B-C). Leukocyte transendothelial migration, NF-kappa B signaling, Chemokine signaling, Th17 differentiation, and Il-17 signaling were represented among differentially expressed genes within each cell type. This data strongly suggests that the *Optn*^−/−^ trigeminal ganglion is subject to severe, chronic, and aberrant inflammation which may be the result of Il-17 signaling, and Cd8 T-cell infiltration.

## Flow cytometry confirms an increase in CD8a-positive T cells and Il-17a expression in T cell populations

To confirm the scRNA-seq results, Cd4 and Cd8 T cell frequencies were characterized for mock infected and HSV-1 infected mice 30 dpi. In the draining lymph nodes (dLNs) there was no difference in the frequency of either Cd4 or Cd8 T cells in the mock groups, but there were significantly more Cd8 T cells in the *Optn*^+/+^ mice (figure 5A-C). Interestingly in the TG of *Optn*^−/−^ mice, there was a near absence of T cells in the mock treatment which increased significantly with infection (figure 5D-F). The *Optn*^+/+^ TG showed similar levels of Cd4 T cells in the *Optn*^+/+^ mock vs HSV-1 infected groups, and similarly to the dLN, the TG showed a decrease in Cd8 T cells, and significantly lower frequencies of Cd8 T cells in the 30 dpi *Optn*^+/+^ vs *Optn*^−/−^ treatments. This supports the higher level of Cd8b1 and Cd8a expression observed in the lymphoid scRNA-seq data.

**Figure 5:**
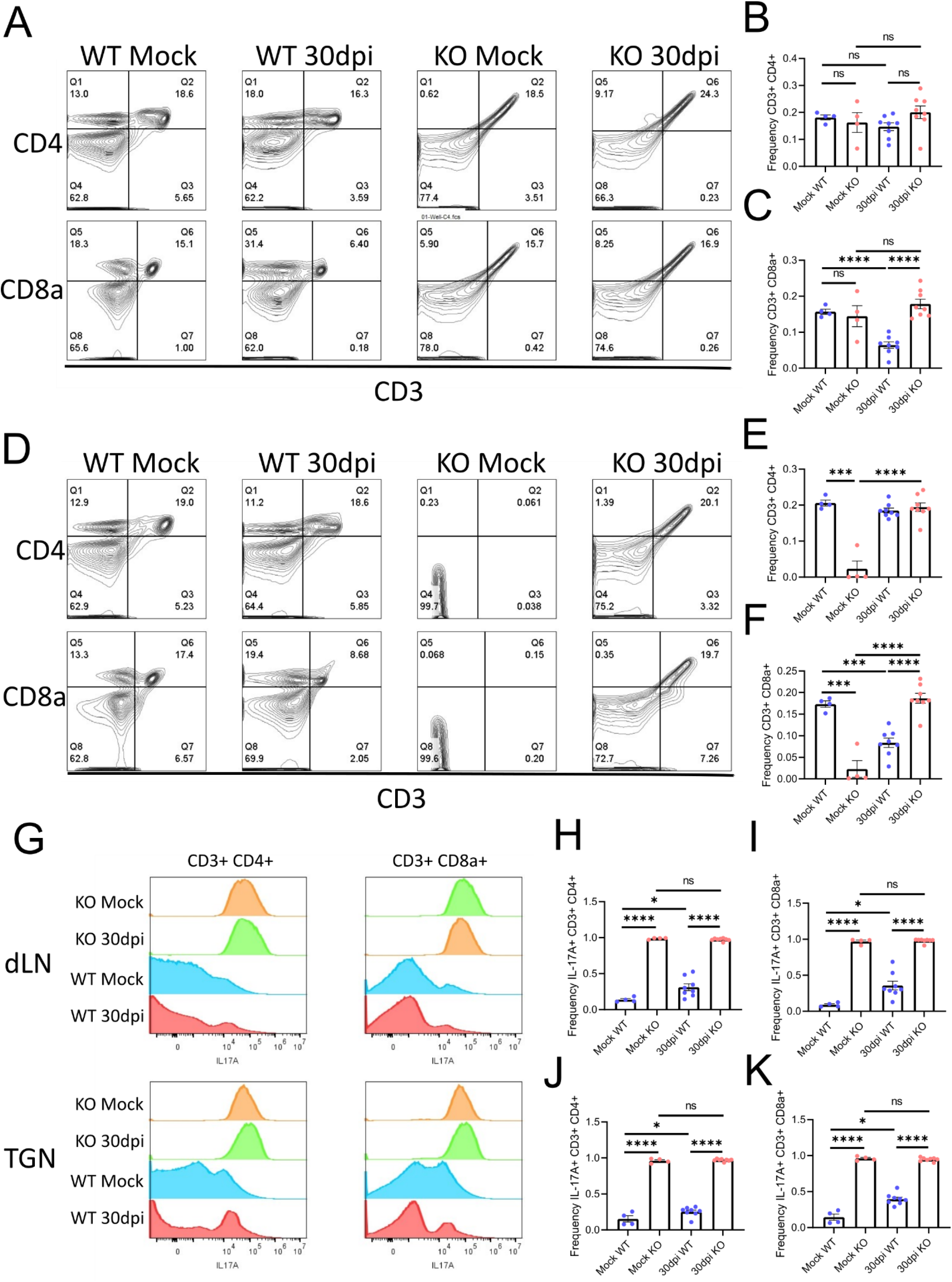
Flow cytometry confirms an increase in CD8a-positive T cells and Il-17a expression in T cell populations. A) Representative contour plots of flow cytometry analysis of CD3+ CD4+ or CD3+ CD8a+ cells in the dLN. B) Quantitation of the frequency for CD3+ CD4+ T-cells in dLNs. C) Quantitation of the frequency for CD3+ CD8a+ T-cells in dLNs. D) Representative contour plots of flow cytometry analysis of CD3+ CD4+ or CD3+ CD8a+ cells in the TGN. E) Quantitation of the frequency for CD3+ CD4+ T-cells in TGN. F) Quantitation of the frequency for CD3+ CD8a+ T-cells in TGN. G) representative histograms showing the distribution of anti-Il-17a intracellular staining in the CD3+ CD4+ or CD3+ CD8a+ T-cells of the dLN (top) or TGN (bottom). H) Quantitation of the frequency for CD3+ CD4+ Il-17a+ T-cells in dLN. I) Quantitation of the frequency for CD3+ CD8a+ Il-17a+ T-cells in dLN. J) Quantitation of the frequency for CD3+ CD4+ Il-17a+ T-cells in TGN. K) Quantitation of the frequency for CD3+ CD8a+ Il-17a+ T-cells in TGN. n >/= 4 mice per group. Student’s t-test was performed for statistical analysis (α= 0.05). *p < 0.05; ***p < 0.001, ****p < 0.0001, ns, not significant.

Further analysis revealed a slight increase in IL-17A positive Cd4 and Cd8 T cells in either the dLN or the TG with infection in the *Optn*^+/+^ mice (figure 5G-K). In sharp contrast, but as predicted by KEGG pathways analysis, *Optn*^−/−^ Cd4 and Cd8 T cells in dLNs and TGs were ubiquitously positive for IL-17A expression. This in in line with the disease presentation of diminished sensitivity and increased keratitis in *Optn*^−/−^ animals.

## *Optn^−/−^* neurons lose neuronal marker expression and undergo demylination despite cell survival

Following detection of a severe T cell driven inflammatory phenotype, we expected that this drives damage to neurons through cell death or demyelination of peripheral neurons in *Optn*^−/−^ mice. Despite the dramatic loss of transcriptionally active neurons and neuronal markers by transcriptomic analysis or staining for SNCG, we did not observe difference in cell death by TUNEL staining (figure 6A-B). There was a difference in fluoromyelin staining at 8 dpi and 30 dpi where *Optn*^−/−^ TGN had diminished staining for myelin compared to *Optn*^+/+^ TGs (figure 6C-D).

**Figure 6:**
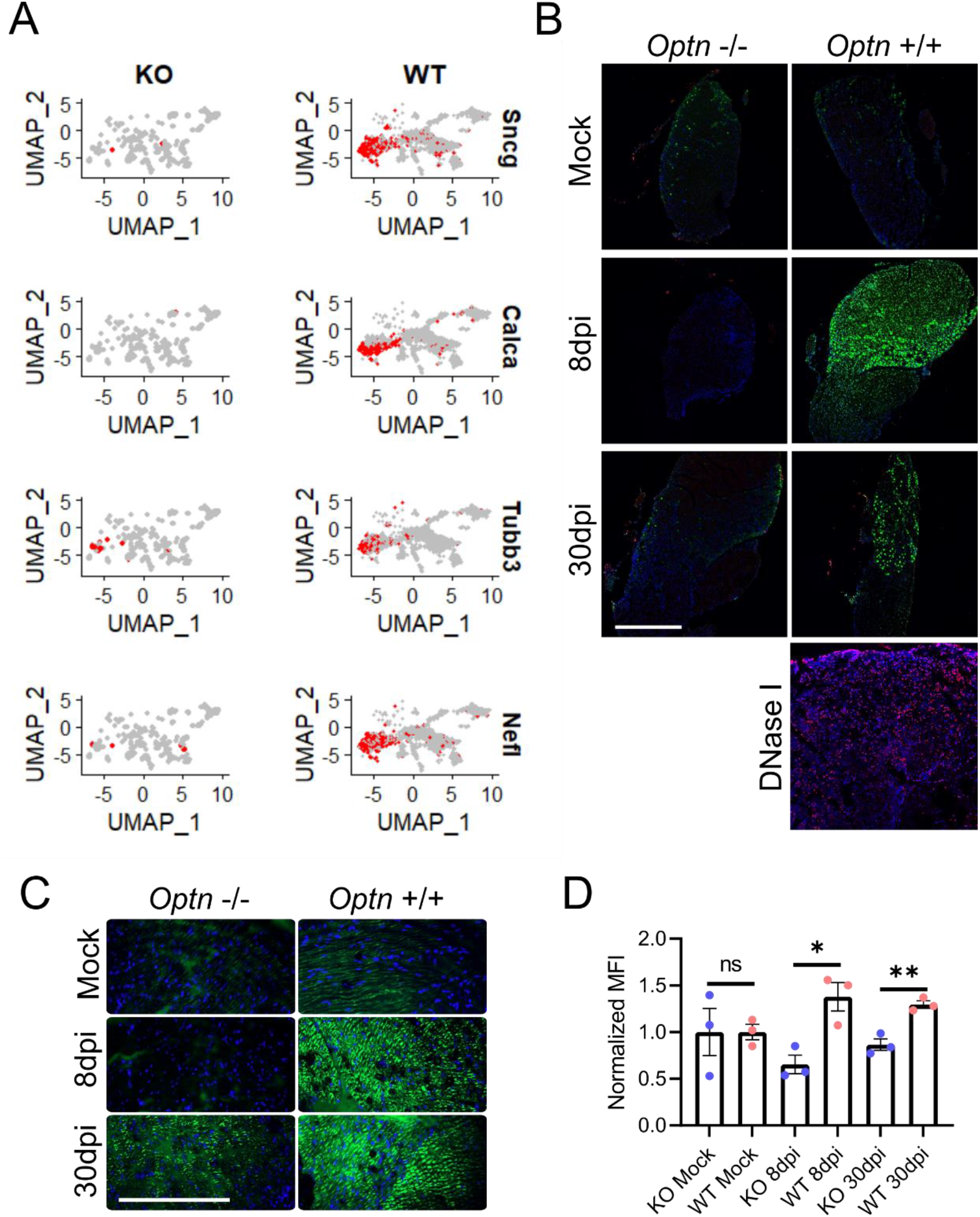
*Optn*−/− neurons lose neuronal marker expression and myelination despite cell survival. A) UMAP analysis for peripheral neuron markers split by genotype reveals distribution of markers across dataset. B) Representative staining for SNCG or TUNEL in mock, 8 dpi, and 30 dpi mouse TGN. Scale bar is 500 μm C) Fluoromyelin staining of mock, 8 dpi, or 30 dpi TGN form WT or KO mice are shown as representative images. D) Quantitation of MFI relative to mock (bottom); n = 3 mice per group. Scale bar is 200 μm Student’s t-test was performed for statistical analysis (α= 0.05). *p < 0.05; **p < 0.01; ns, not significant.

## Conclusion

We highlight the importance of OPTN in protection against herpes keratitis pathologies. Mice corneally infected with HSV-1 showed similar level of viral replication in corneal tissue and in ocular washes, but despite this the *Optn*^−/−^ mice demonstrated a rapid loss of corneal and whisker sensitivity along with severe opacification of the cornea. Clinically, ocular HSV-1 can be treated with acyclovir, which is highly effective in halting non-resistant strain of HSV-1 ^6^. Patients must then take corticosteroids for an extended period to suppress the HSV-1 associated inflammation ^6^. Our *Optn*^−/−^ mouse model of ocular HSV-1 infection recapitulates the clinical consequences of untreated herpes keratitis as the *Optn*^−/−^ mice show signs of corneal scaring and lost sensation. Often asymptomatic individuals can shed HSV-1 without any sign of disease, and in our model the wildtype animals represented a group that shed HSV-1 despite having only mild disease ^7^.

scRNA-seq revealed that OPTN-deficiency leads to severe long-term loss of neuronal function in the TG, though the exact mechanism remains unclear. Despite the loss of transcription in neurons, they are not undergoing cell death. We were able to confirm the expression pattern of our top identified neuronal biomarker, SNCG, hereby increasing confidence in the scRNA-seq analysis. DGE and KEGG pathways enrichment analysis revealed upregulation of many immune-signaling pathways in *Optn*^−/−^ Schwann cells, satellite glial cells, and macrophages. While it was expected to see differences in pathways that OPTN has been previous implicated in, we were surprised to have seen representation of the terms T cell receptor signaling pathway, Th1 and Th2 cell differentiation, IL-17 signaling pathway, and Th17 cell differentiation across multiple cell types. We previously highlighted that OPTN is required for effective recruitment of CD4 and CD8 T-lymphocytes to the CNS for protection against herpes encephalitis. The upregulation of these pathways provides a research direction to further investigate the role of T-lymphocyte mediated anti-HSV-1 immunity in the peripheral nervous system.

It is previous proposed that CD8 T-lymphocytes are indispensable in suppressing reactivation of HSV-1, however there is only a single study of the role of a Th17 response against HSV-1 infection where infection of IL-17R knockout mice resulted in diminished keratitis severity ^8, 9^. In another study, it has been shown that IL-17 treatment of Schwann cells results in the decreased ability of these cells to myelinate axons ^10^. Here we show that OPTN suppresses excessive IL-17 expression in T lymphocytes and subsequent demyelination in the TG. Consistent with demyelination the *Optn*^−/−^ mice lose whisker and corneal sensitivity.

## Materials

### Mice

The C57/b6J *Optn*^−/−^ mouse model used in this study are outline elsewhere in ^5, 11^. Wildtype C57/b6J mice were purchased from Jackson Laboratories, Bar Harbor, ME, USA. Male mice were used in this study and all mice were 8-12 weeks old at the beginning of each experiment.

### Viruses

McKrae strain Herpes simplex virus type-1 used throughout this study was provided by Dr. Patricia Spear (Northwestern University).

### Antibodies and stains

The following antibodies and stains were used in this study: Anti-gamma Synuclein/SNCG antibody (ab55424) (Abcam), In Situ Cell Death Detection Kit, TMR red (Roche), DAPI (Sigma), Goat anti-Rabbit IgG (H+L) Cross-Adsorbed Secondary Antibody, Alexa Fluor 488 (Invitrogen), FluoroMyelin Green Fluorescent Myelin Stain (Invitrogen), Brilliant Violet 605^™^ anti-mouse IL-17A Antibody clone [TC11-18H10.1], Alexa Fluor^®^ 700 anti-mouse CD3 Antibody [17A2] (BioLegend), APC/Cyanine7 anti-mouse CD8a Antibody [53-6.7] (BioLegend), Brilliant Violet 605^™^ anti-mouse IL-17A Antibody [TC11-18H10.1] (BioLegend), PE Anti-Mouse CD4 [RM4-5] (BioLegend)

### Software

The software used in this study includes Partek Flow (Partek, Inc), GraphPad Prism 9 (GraphPad Software, San Diego, CA), ZEN 3.1 Blue edition (Carl Zeiss, Jena, Germany), RStudio (RStudio, PBC) with R 4.1.1, and ImageJ.

## Methods

### Infections

The corneal scarification method was used for infection of mice as published previously ^5^. In brief mice were anesthetized using ketamine (100 mg/kg) and xylazine (5 mg/kg). Proparacaine hydrochloride (1% ophthalmic topical solution) was applied to the corneal surface while the mice were unconscious. Corneal epithelial debridement was performed with a 30-gauge needle, then 1×10^5^ PFU HSV-1 in a 5 μl volume of PBS was immediately applied to the cornea.

### Flow Cytometry

Tissues were dissociated into single cell suspensions immediately following dissection. dLNs or TGs were digested in 5 mg/mL collagenase D (Roche) at 37 °C for 2 hours with constant agitation. Samples were then triturated using a pipette, filtered through a 100 μm strainer then fixed for 15 minutes in 4% paraformaldehyde. One million cells were then suspended in Flow assisted cell sorting (FACS) buffer (PBS, 1% BSA) and samples were loaded into a 96-well round bottom cell dish. For staining, cells were blocked and permeabilized with PBS, 10% BSA, 0.1% Triton-X and TruStain FcX (BioLegend) according to manufacturer’s protocol. After blocking, 2 rounds of centrifugations and washes in 100 μL FACS buffer were performed. An antibody cocktail was added to cells according to manufacturer’s recommendations and incubated on ice in darkness for 30 minutes. Two more rounds of centrifugations and washes in 100 μL FACS buffer were performed before samples were submitted to the University of Illinois at Chicago flow cytometry core to be analyzed on a BD CytoFlex flow cytometer. Flow cytometry data was analyzed using FlowJo software (Tree Star Inc.).

### DROPseq generation of single cell libraries

Generation of single cell libraries was performed exactly as outlined by Macosko et al., 2015 ^12^. In summary TGs were dissociated into single cell suspensions by digestion with the Papain Dissociation System (Worthington Biochemical Corporation, Lakewood, NJ) according to manufacturer’s protocol. Cells were mixed into droplets with barcoded beads using a microfluidic chip. Droplets were disrupted and cDNA synthesis and PCR were performed to amplify libraries following library preparation using the Illumina Nextera XT kit. Sequencing was performed by the University of Illinois at Chicago DNA services core facility on an Illumina NextSeq.

### scRNA-seq analysis

Quality control, alignment to the mouse genome, and generation of the cell expression matrix was performed on Parktek servers using Partek Flow software according to the developer’s manual. The cell expression matrix was then downloaded and read into RStudio as a SeuratObject for analysis with the Seurat R package according to the Satija Laboratory vignettes. Following generation of a SeuratObject for the WT and KO data, the ‘FindVariableFeatures’ function was used to select the top 2000 variables features. The WT and KO data were then combined by using the ‘FindIntegrationAnchors’ for the first 20 dimensions followed by ‘IntergrateData’ using the integration anchors. We then used the ‘ScaleData’ functions on the newly combined SeuratObject and ran the ‘RunPCA’ function with “features = VariableFeatures”. We decided to use the first 15 principal components and generated clusters using the ‘FindNeighbors’ function with principal components 1 through 15 followed by ‘FindClusters’ with “resolution = 0.15”. For each cluster ‘FindConservedMarkers’ was used to identify the biomarkers for each cluster. These biomarkers were used to perform literature-based annotation for cluster celltypes. We identified contaminating cell types from the pituitary gland, and these were removed from analysis. To obtain DGE data, the ‘FindMarkers’ function was used with the negative binomial model. The PathfindR R package was used according to the publisher’s vignette to identify the enriched KEGG pathways for each cluster using the output of the DGE analyses from Seurat.

### Histology

Method for sample preparation for histological analysis is outline elsewhere^5^. Following euthanasia and dissection of mice, tissue was fixed overnight in 4% PFA at 4 °C. After fixation tissue was cryoprotected in a 30% sucrose in PBS solution at 4 °C until the tissue sank to the bottom of the tube. Tissue was then embedded in OCT and frozen on crushed dry ice. 10 μm sections were cut and mounted onto Superfrost Plus microscope slides (Fisher Scientific) using a NX10 Cryostat microtome. Hematoxylin and eosin staining was performed as described elsewhere^13^. For myelin staining, slides were permeabilized with PBT (PBS with 0.1% Triton X-100) with 10% BSA for 1 hour at room temperature. Slides were washed three times with PBT before staining with FluoroMyelin^™^ Green Fluorescent Myelin Stain (Invitrogen) at 1:300 dilution and DAPI for 2 hours at room temperature in darkness. Slides were washed 3 times in PBT before mounting with Vectashield mounting solution (Vector laboratories) and cover glass.

### Immunofluorescence Staining

For immunofluorescence staining the same preparation as above was followed through the permeabilization and blocking steps. The primary antibody and TUNEL reaction mixture were added to slides and incubated at 37 °C for 1 hour. Slides were washed 3 times in PBS then the secondary antibody and DAPI were applied in PBS for 1 hour at room temperature. Slides were washed 3 times in PBS before mounting with Vectashield mounting solution (Vector laboratories) and cover glass.

### Plaque Assay

Titration of HSV-1 was performed by plaque assay as outlined elsewhere ^5^. For ocular washes, samples were collected by pipetting a 10 μL droplet of PBS onto the corneal surface, then drawing the liquid back up into the tip and dispelling the sample into 300 μL of Opti-MEM. The sample was used for the same plaque assay protocol referenced.

### Statistical analysis

All statistical analysis and graph making was carried out in GraphPad Prism software, except for flow cytometry histograms and dot plots produced in FlowJo software. Error bars represent ± SEM of at least three independent measurements (n = 3). Asterisks denote a significant difference, as determined by two-tailed unpaired Student’s t test, Mann-Whitney U test, or Logrank test: *p < 0.05; **p < 0.01; ***p < 0.001; ns, not significant, or two-way ANOVA; ****p < 0.0001.

